# Homogenic endothelium-independent hematopoiesis ensures the abundance of erythrocytes and macrophages during zebrafish development

**DOI:** 10.1101/2023.02.20.529211

**Authors:** Ramy Elsaid, Aya Mikdache, Keinis Quintero Castillo, Yazan Salloum, Patricia Diabangouaya, Gwendoline Gros, Carmen G. Feijoo, Pedro P. Hernández

## Abstract

In all organisms studied, from flies to humans, blood cells emerge in several sequential waves and from distinct hematopoietic origins. However, the relative contribution of these ontogenetically distinct hematopoietic waves to embryonic blood lineages and to tissue regeneration during development is yet elusive. Here, using a lineage-specific ‘switch and trace’ strategy in the zebrafish embryo, we report that the definitive hematopoietic progeny barely contributes to erythrocytes and macrophages during early development. Lineage tracing further show that ontogenetically distinct macrophages exhibit differential recruitment to the site of injury based on the developmental stage of the organism. We further demonstrate that primitive macrophages can solely maintain tissue regeneration during early larval developmental stages after selective ablation of definitive macrophages. Our findings highlight that the sequential emergence of hematopoietic waves in embryos ensures the abundance of blood cells required for tissue homeostasis and integrity during development.

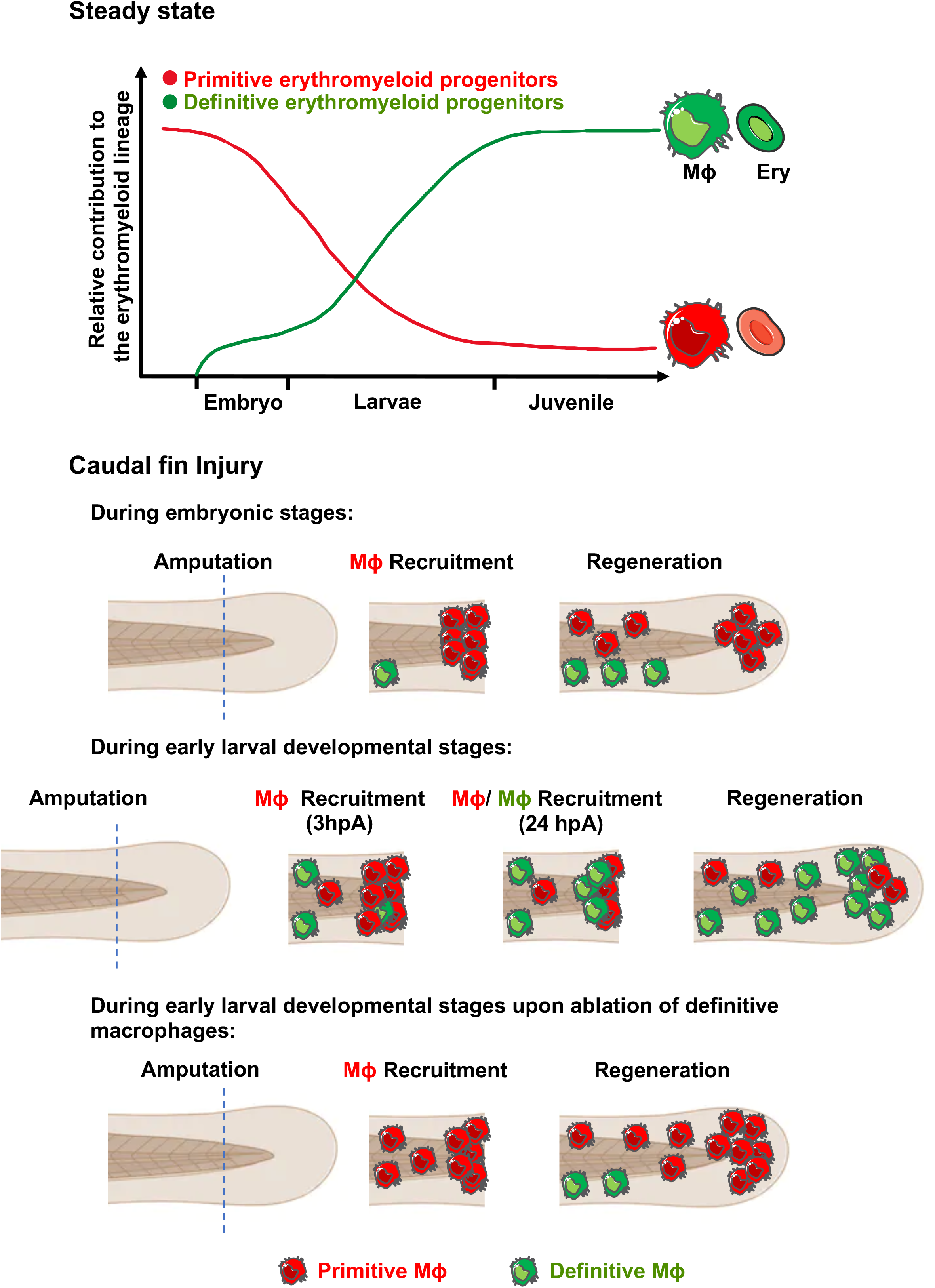

## Introduction

Hematopoiesis is a complex biological process by which all mature blood lineages are generated. In mammals, hematopoiesis originates from distinct blood progenitors that emerge during development through an endothelial-to-hematopoietic transition mechanism (EHT) ^1–3^, resulting in a layered organization of the immune system ^4^. As in mammals, zebrafish hematopoiesis also consists of multiple waves that emerge at distinct anatomical locations ^4^. In the zebrafish embryo, primitive hematopoiesis emerges intra-embryonically directly from mesoderm around 11 hours post-fertilization (hpf) in the rostral blood island (RBI) and the posterior lateral mesoderm (PLM) and generates primitive erythrocytes and myeloid cells ^4,5^. Other definitive or endothelial-derived hematopoietic waves are also produced in the developing organism ^4^. One of them is a transient wave that emerges from the posterior blood island (PBI) at 24-30 hpf via EHT and gives rise to erythro-myeloid progenitors (EMPs) like those in mammals ^6–8^. At 36 hpf, other waves of HSCs and HSC-independent progenitors emerge from the dorsal aorta (DA) via EHT ^1,3^. These DA-derived hematopoietic progenitors then migrate to the caudal hematopoietic tissue (CHT), the fetal liver counterpart in zebrafish^9^. Therefore, due to its similarity to mammalian hematopoiesis, the zebrafish is an excellent model to study the ontogeny of different hematopoietic waves during development.

Although it is well characterized that blood cells are produced in sequential and overlapping waves ^4^, little is known about their contribution to different hematopoietic lineages during development. Recently, independent reports support the notion that definitive hematopoiesis sustains embryonic blood lineages ^8,10–12^. However, primitive hematopoietic progenitors give rise also to the erythromyeloid lineage and it is yet unknown to what extent these progenitors contribute to embryonic blood lineages. This is mainly due to the lack of specific markers for primitive hematopoietic progenitors in mammals ^4,7,13,14^.

The difficulty in accurately tracing the output of each wave in mammalian animal models has hindered the full understanding of immune cell ontogeny. In this study, we used the unique strengths of the zebrafish embryo to perform temporarily resolved lineage tracing combined with live imaging to investigate the embryonic erythromyeloid lineage ontogeny *in vivo*. We unveiled that embryonic erythromyeloid lineages originate from primitive hematopoietic progenitors with a delayed contribution from definitive hematopoietic waves to the erythromyeloid lineage till the larval developmental stage. We further showed that primitive macrophages were recruited to the damage site earlier than their definitive counterparts. Our results also suggest that primitive macrophages maintain tissue regeneration during early larval developmental stages after selective ablation of definitive macrophages. Combined, our study reveals that the sequential emergence of hematopoietic waves ensures the abundance of macrophages and erythrocytes required for tissue homeostasis and integrity during development.

## Results

### Primitive hematopoietic progenitors sustain erythrocytes during embryonic development

As all definitive hematopoietic progenitors originate from hemogenic endothelium (HE) as early as 24 hpf ^1,6,8,10,15,16^, while primitive progenitors originate directly from mesoderm in zebrafish^1,3,17,18^, we set up a strategy to distinguish between primitive and definitive hematopoietic progenitors based on their tissue of origin. We used the endothelial-specific tamoxifen-inducible transgenic line *Tg(fli1a:CreERT2)* ^19^ in combination with different hematopoietic lineage-specific switch and trace lines^20^. Our inducible labeling strategy at 24 hpf will label exclusively definitive hematopoietic progenitors as they emerge from the HE without labeling primitive hematopoietic progenitors. To assess the efficiency of our labeling strategy, we combined the *Tg(fli1a:CreERT2)* line with two different independent transgenic lines, the lymphocyte-specific *Tg(lck:loxP-DsRedx-loxP-GFP)* ^8,21^ line (supplemental Figure 1A) and the *Tg(coro1a:loxP-DsRedx-loxP-GFP)* line which labels all leukocytes^13,22^ (supplemental Figure 1D). We found that 80% of thymocytes in the thymus, which have only a definitive hematopoietic origin ^8,23^, were labelled using either the *Tg(lck:loxP-DsRedx-loxP-GFP)* (supplemental Figure 1B-C) or the *Tg(coro1a:loxP-DsRedx-loxP-GFP)* lines (supplemental Figure 1E-F), indicating the high labelling efficiency of our system.

To determine the origin of erythrocytes during embryonic development and early larval stages, we combined the *Tg(fli1a:CreERT2)* line with the erythrocyte-specific *Tg(α/β_a2_-globin:loxP-DsRedx-loxP-GFP)* line (referred to hereafter as globin:switch) ^8,18^. In double transgenic zebrafish, 4-OH-tamoxifen (4-OHT) induces cre recombination and removes the DsRed cassette, leading to permanent GFP expression in *fli1a*^+^-derived erythroid progeny (Figure 1A). To determine the contribution of definitive hematopoietic progenitors to erythropoiesis during early stages of development, we exposed embryos to either 4-OHT or ethanol (EtOH) starting at 24 hpf to label the aortic-endothelium-derived definitive hematopoietic progenitors ^1,8,15^, and then monitored GFP^+^ definitive erythrocytes via live imaging (Figure 1B and supplemental video 1). Primitive hematopoietic progenitors robustly generated erythrocytes from early embryonic stages (Figure 1B). In contrast, GFP^+^ definitive erythrocytes started to emerge after 2 dpf, accumulated in the CHT region starting at 4 dpf (Figure 1C) and their contribution to the erythrocyte pool was not appreciable until later larval developmental stages (∼12 dpf, Figure 1C-F).

**Figure 1:**
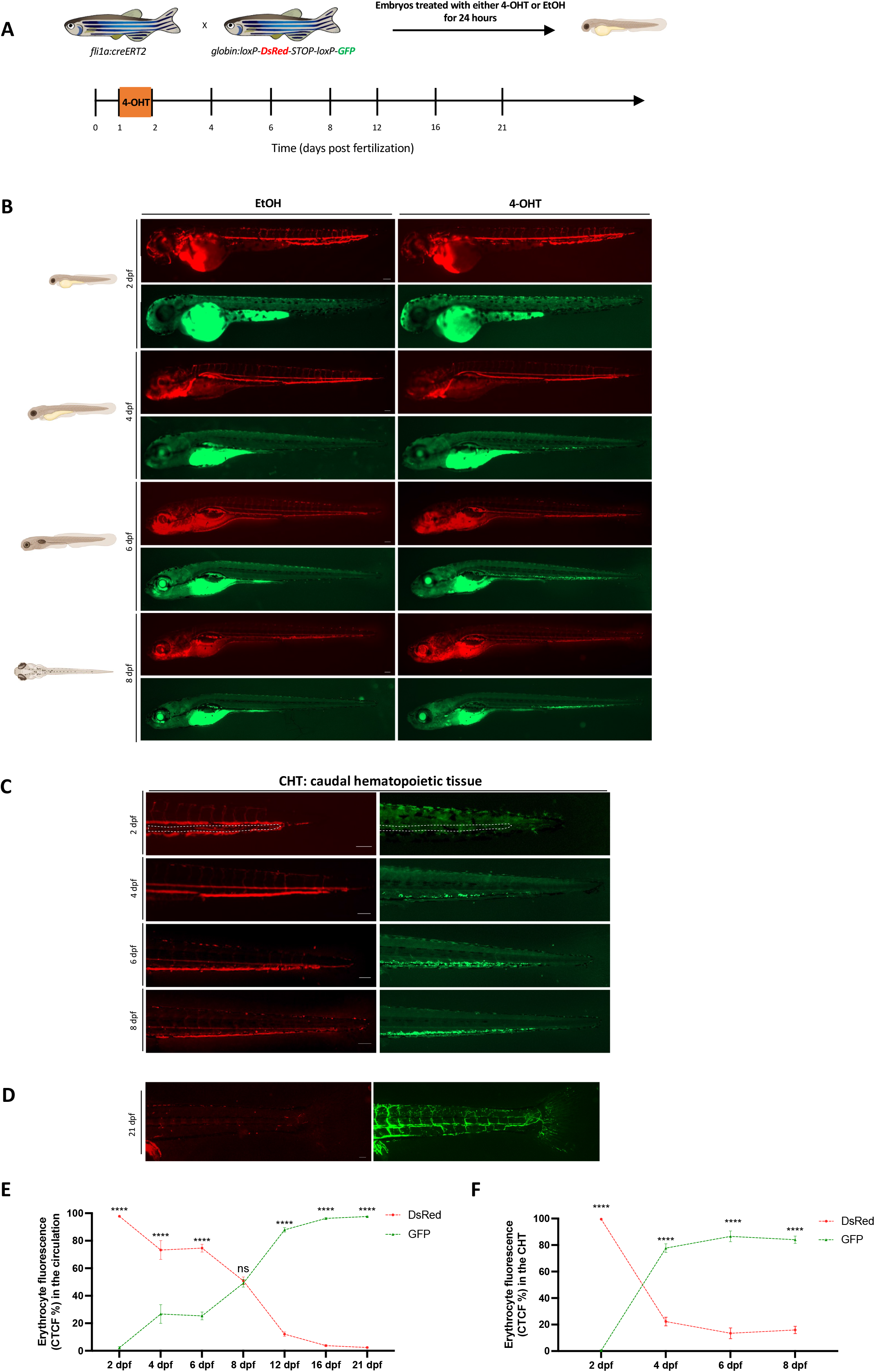
Tracing the contribution of definitive hematopoietic waves to the erythroid lineage in the embryoand early larvae. (A) Scheme of the 4-OHT-inducible transgenic lines used to assess the contribution of definitive hematopoietic waves to the erythroid lineage. (B) Fluorescent images of EtOH non-switched controls (left) and 4-OHT-induced (right) *Tg(fli1a:creERT2;globin:Switch)* embryos and larvae (2 dpf-8 dpf). Scale bars, 100 µm. Quantification of erythrocytes in circulation are shown in (E). (C) Fluorescent images of 4-OHT-induced *Tg(fli1a:creERT2;globin:Switch)* embryos and larvae (2-8 dpf) in the CHT region. Non-switched primitive erythrocytes (left) and definitive erythrocytes (right). Scale bars, 100 µm. quantification of erythrocytes in the CHT are shown in (F). (D) Fluorescent images of 4-OHT-induced *Tg(fli1a:creERT2;globin:Switch)* (21 dpf) in the circulation. Scale bar,100 µm. Non-switched primitive erythrocytes (left) and definitive erythrocytes (right). Quantification of erythrocytes in the circulation are shown in (E). (E) Quantification of DsRed and GFP fluorescence intensity percentage in the circulation over a time course of 2-21 dpf. (2 dpf n=6; 4 dpf n=4; 6 dpf n=5; 8 dpf n=4; 12 dpf; 16 dpf n=6 and 21 dpf n=5). Mean ± SEM of the DsRed^+^ and GFP^+^ corrected total cell fluorescence (CTCF) percentage at each time point is shown. Two-way ANOVA with Sidak’s multiple comparison was used for this analysis. ^∗∗∗∗^p ≤ 0.0001. (F) Quantification of DsRed and GFP fluorescence intensity percentage in the CHT over a time course of 2-8 dpf. (2 dpf n=6; 4 dpf n=4; 6 dpf n=5 and 8 dpf n=4). Mean ± SEM of the DsRed^+^ and GFP^+^ corrected total cell fluorescence (CTCF) percentage at each time point is shown. Two-way ANOVA with Sidak’s multiple comparison was used for this analysis. ^∗∗∗∗^p ≤ 0.0001.

These results indicate that erythrocytes emerge in a layered organization during embryonic and early larval stages as in mammals ^24^, and that primitive hematopoietic progenitors generate a sufficient number of erythrocytes to sustain embryonic survival and tissue homeostasis ^25^.

### Embryonic macrophages originate exclusively from primitive hematopoietic progenitors

To assess the contribution of definitive hematopoietic progenitors to the embryonic macrophage pool, we marked the latter by combining the *Tg(fli1a:CreERT2)* line with the macrophage-specific *Tg(mpeg1:LoxP-DsRedx-LoxP-GFP-NTR)* line (referred to hereafter as mpeg1:switch) ^26,27^. As aforementioned, 4-OHT-induced cre recombination leads to permanent GFP expression in *fli1a^+^*-derived macrophages progeny (Figure 2A). Using this system, embryonic microglia cells, which are known to be of primitive hematopoietic origin in zebrafish ^7,13^, were not labelled, indicating the precision of this labelling system (supplemental figure 2A-B). While primitive macrophages contributed robustly to the embryonic macrophage pool, we found that definitive hematopoietic progenitors minimally contribute to macrophages during early developmental stages, akin to erythrocytes ontogeny (Figure 2B). We showed further that GFP^+^ definitive macrophages started to emerge after 2 dpf and gradually increased in numbers in the CHT region (Figure 2B and E) with a delayed modest contribution to the macrophage pool in the periphery starting after 4 dpf (Figure 2B-D). GFP^+^ definitive macrophages contributed robustly to peripheral macrophages by 16 dpf, suggesting a distinct differentiation kinetics of primitive and definitive macrophages throughout development (Figure 2C).

**Figure 2:**
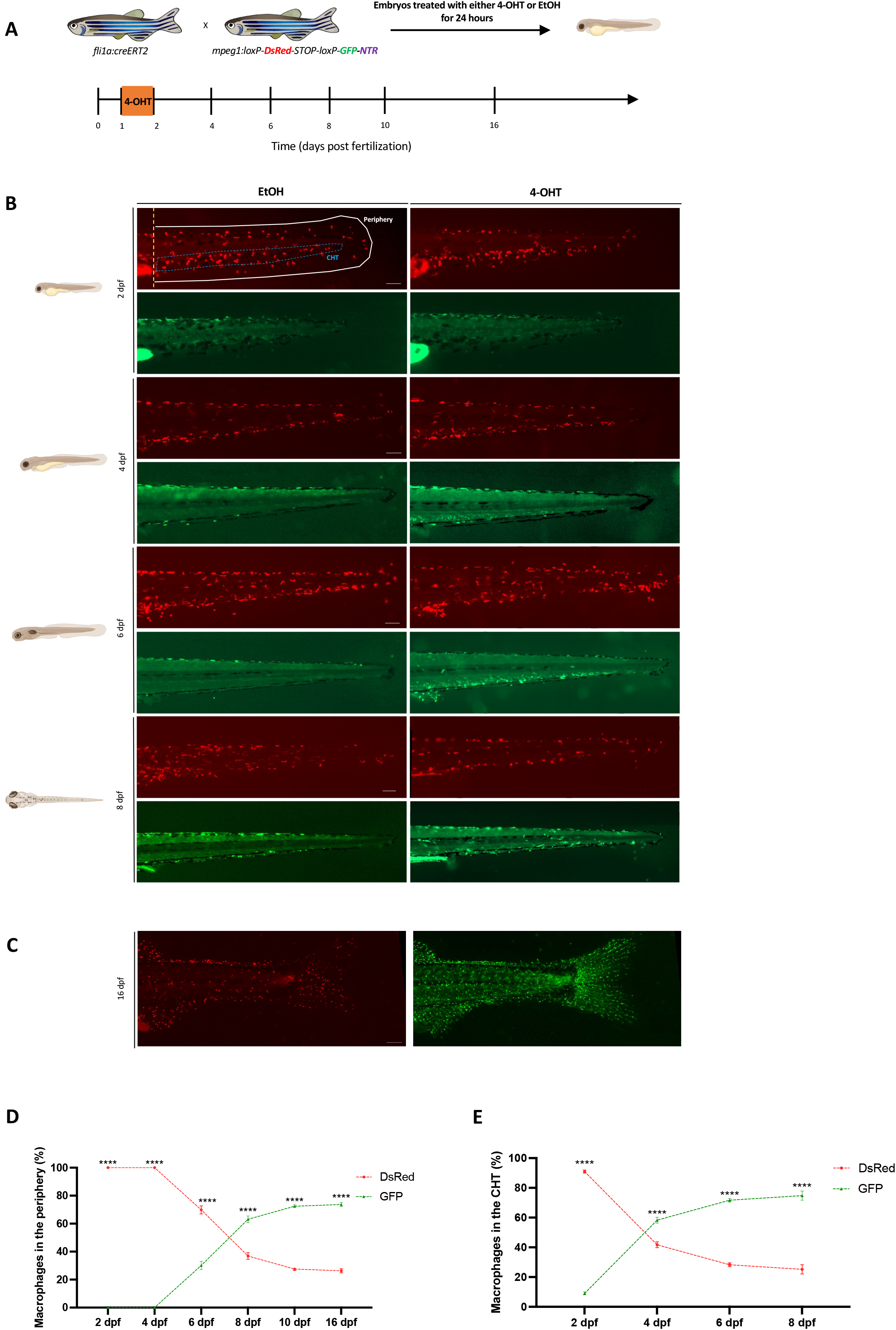
Tracing the contribution of definitive hematopoietic waves to macrophages in the embryo and early larvae. (A) Scheme of the 4-OHT-inducible transgenic lines used to assess the contribution of definitive hematopoietic waves to the macrophage lineage. (B) Fluorescent images of EtOH non-switched controls (left) and 4-OHT-induced (right) *Tg(fli1a:creERT2;mpeg1:Switch)* embryos and larvae (2-8 dpf). Macrophages were considered as peripheral macrophages (white area) or CHT-resident macrophages (blue area). Scale bars, 100 µm. Quantification of macrophages in the periphery and the CHT are shown in (D) and (E) respectively. (C) Fluorescent images of 4-OHT-induced *Tg(fli1a:creERT2;mpeg1:Switch)* (16 dpf) in the periphery. Scale bar, 500 µm. Non-switched primitive macrophages (left) and definitive macrophages (right). Quantification of macrophages in the periphery are shown in (D). (D) Quantification of DsRed+ and GFP+ macrophages in the periphery over a time course of 2-16 dpf. (2 dpf n=6; 4 dpf n=11; 6 dpf n=7; 8 dpf n=4; 10 dpf n=5 and 16 dpf n=5). Mean ± SEM of the DsRed^+^ and GFP^+^ macrophage number at each time point is shown. Two-way ANOVA with Sidak’s multiple comparison was used for this analysis. ^∗∗∗∗^p ≤ 0.0001. (E) Quantification of DsRed+ and GFP+ macrophages in the CHT over a time course of 2-8dpf. (2 dpf n=6; 4 dpf n=11; 6 dpf n=7 and 8 dpf n=4). Mean ± SEM of the DsRed^+^ and GFP^+^ macrophage numberat each time point is shown. Two-way ANOVA with Sidak’s multiple comparison was used for this analysis. ^∗∗∗∗^p ≤ 0.0001.

Altogether, our lineage tracing experiments show that aortic endothelium derived hematopoietic progenitors are barely contributing to embryonic erythroid and myeloid lineages (Figures 1 and 2).

### Ontogenically distinct macrophages are differentially recruited to the site of injury

It has been recently reported that distinct macrophage subpopulations play different roles to promote tail regeneration after amputation ^28,29^. However, whether ontogenetically distinct macrophages display different functions and recruitment behaviors to the site of injury remains poorly understood. Therefore, we sought to analyze if macrophages of distinct origins are recruited in a differential manner to the damage site at different developmental stages. Using the same labelling strategy (Figure 2A), we analyzed macrophages recruitment to the damaged site in embryos (2 dpf) and larvae (5 dpf) since our lineage tracing experiments showed a different abundance of ontogenetically distinct macrophages at these developmental stages (Figure 2). In embryos, we observed that primitive macrophages were recruited to the site of injury at 24 hours post amputation (hpa) and 48 hpa (Figure 3A-B and supplemental Figure 3A). We then analyzed macrophage recruitment to the damage site in larvae, where both primitive and definitive macrophage populations coexist in the periphery.

**Figure 3:**
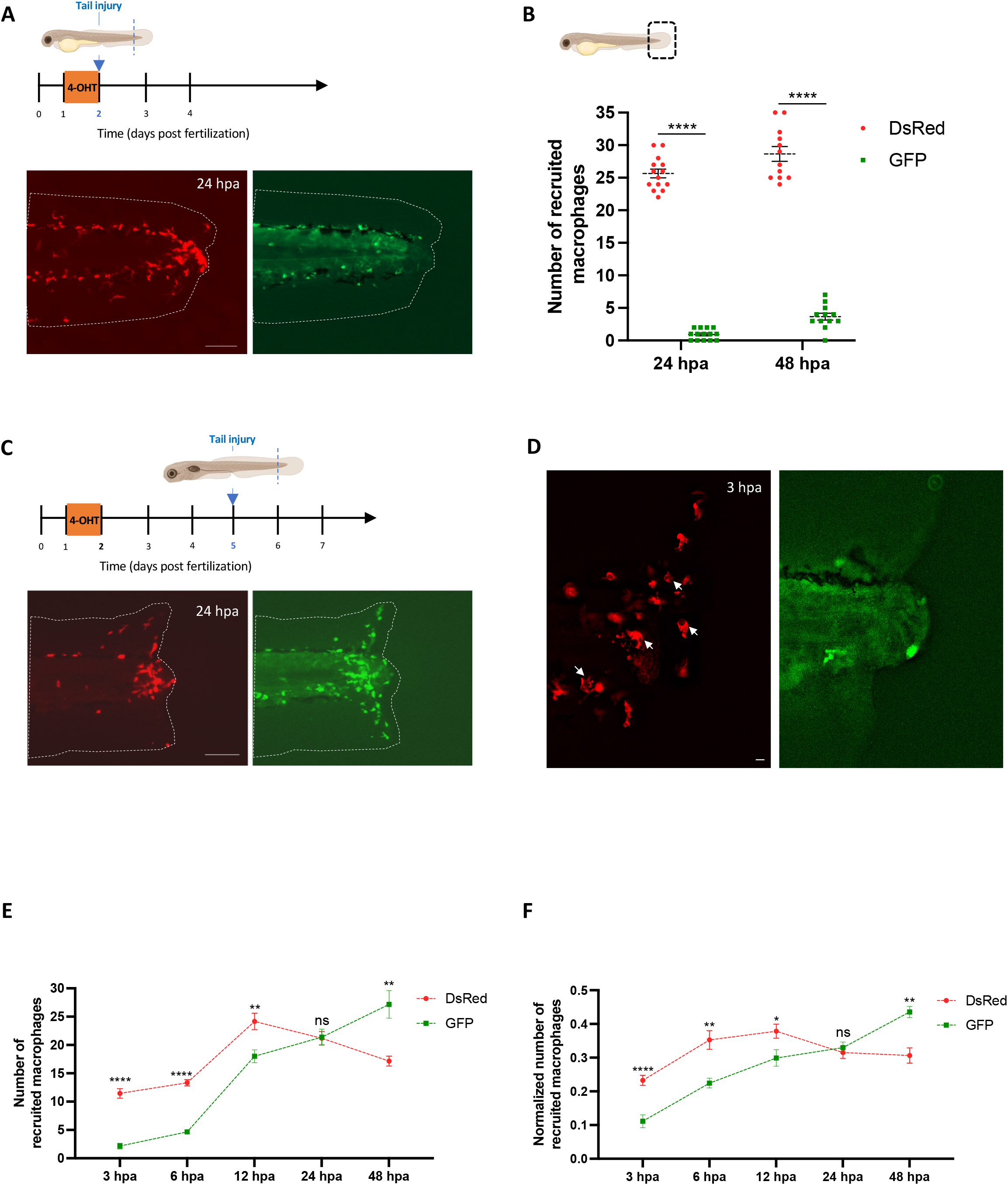
Ontogenetically distinct macrophages exhibit different migration behavior and recruitment aftertissue injury at early developmental stages. (A) Diagram showing the amputation plan, tail fin of *Tg(fli1a:creERT2;mpeg1:Switch)* were amputated at 2 dpf and macrophages recruitment to the site of injury was analyzed. Representative images are shown at 24 hpa.Scale bar: 100µm. (B) Diagram showing the counting region in the larvae. Quantification of DsRed+ and GFP+ macrophages at the site of injury at 24 hpa (n=14) and 48 hpa (n=12). Mean ± SEM of the DsRed^+^ and GFP^+^ macrophage number at each time point is shown. ^∗∗∗∗^p ≤ 0.0001. (C) Diagram showing the amputation plan, tail fin of *Tg(fli1a:creERT2;mpeg1:Switch)* were amputated at 5 dpf and macrophage numbers at the site of injury was analyzed. Representative images are shown at 24 hpa. Scale bar: 100µm. (D) Tail fin of *Tg(fli1a:creERT2;mpeg1:Switch)* were amputated at 5 dpf and macrophage numbers at the site of injury was analyzed. Representative images are shown at 3 hpa. White arrows indicate the primitive macrophages with big vacuoles. Scale bar: 10µm. (E) Quantification of DsRed+ and GFP+ macrophages at the site of injury at 3 hpa (n=13); 6 hpa (n=6); 12 hpa (n=6); 24 hpa (n=10) and 48 hpa (n=6). Mean ± SEM of the DsRed^+^ and GFP^+^ macrophage number at each time point is shown. ns, p>0.05; ^∗∗^p ≤ 0.01; ^∗∗∗∗^p ≤ 0.0001. (F) Quantification of DsRed+ and GFP+ macrophages at the site of injury at 3 hpa (n=13); 6 hpa (n=6); 12 hpa (n=6); 24 hpa (n=10) and 48 hpa (n=6). Mean ± SEM of the DsRed^+^ and GFP^+^ macrophage numbers at each time point is shown. Quantification was normalized by the number of total macrophages in the tail of the respective larvae (the sum of peripheral, CHT and recruited macrophages of distinct origins). ns, p>0.05; ^∗^p ≤ 0.05; ^∗∗^p ≤ 0.01; ^∗∗∗∗^p ≤ 0.0001.

We did not observe differences in the recruitment of both primitive and definitive macrophages to the damage site 24 hpa with a slight increase in the number of definitive macrophages 48 hpa (Figure 3C, E and supplemental Figure 3B). Altogether, these results suggest that primitive macrophages can alone modulate tail fin regeneration during embryonic stages while both macrophage populations are recruited to the damaged site in the larvae.

Previous reports showed that the first recruited macrophages play an important role in proper tail fin regeneration ^29–31^, therefore, we further analyzed which macrophage population is recruited first to the site of injury. We found that primitive macrophages are recruited earlier than their definitive counterparts showed by the reduced number of definitive macrophages recruited to the site of injury from 3 to 12 hpa (Figure 3E-F). In addition, primitive macrophages manifested sings of active phagocytosis activity as early as 3 hpa evidenced by the appearance of vacuoles in their cell bodies (Figure 3D). Since our observation of a delay in the recruitment of definitive macrophages could be the result of differences in the number of primitive and definitive macrophages in the periphery at the time of injury, we attempted to normalize their numbers. Data was normalized by dividing the number of each recruited macrophage subpopulation by their total number in the tail of the same larvae ^29^. After normalization, we still observed that primitive macrophages are recruited earlier compared to their definitive counterparts with a reduced recruitment of definitive macrophages at 3, 6 and 12 hpa, when compared to their primitive counterparts (Figure 3F). These data indicate that primitive macrophages, as the first arrivals to the damage site, may have a specific role in the regeneration process after amputation during development.

### Selective ablation of definitive macrophages does not impair tail fin regeneration during early larval developmental stages

To evaluate the contribution of ontogenetically distinct macrophages to tail fin regeneration, we used the mpeg1 switch line *Tg(mpeg1:loxP-dsRed-loxP-eGFP-NTR)* that allows selective ablation of macrophages based on their origin. In this line, the expression of bacterial nitroreductase (NTR) is under the control of the *mpeg1* promoter ^27^. Thus, the NTR will be expressed exclusively in the traced macrophages, thereby, traced macrophages could be ablated by metronidazole (MTZ) treatment ^32^ (Figure 4A). To assess the efficiency of the NTR-MTZ system, we quantified the number of primitive and definitive macrophages in the tail after treatment with either DMSO or MTZ for 48 hours from 4 to 6 dpf. We found that while primitive macrophages were not affected by the MTZ treatment, definitive macrophages were significantly reduced after 48 hours of MTZ treatment (Supplemental Figure 3C-D).

**Figure 4:**
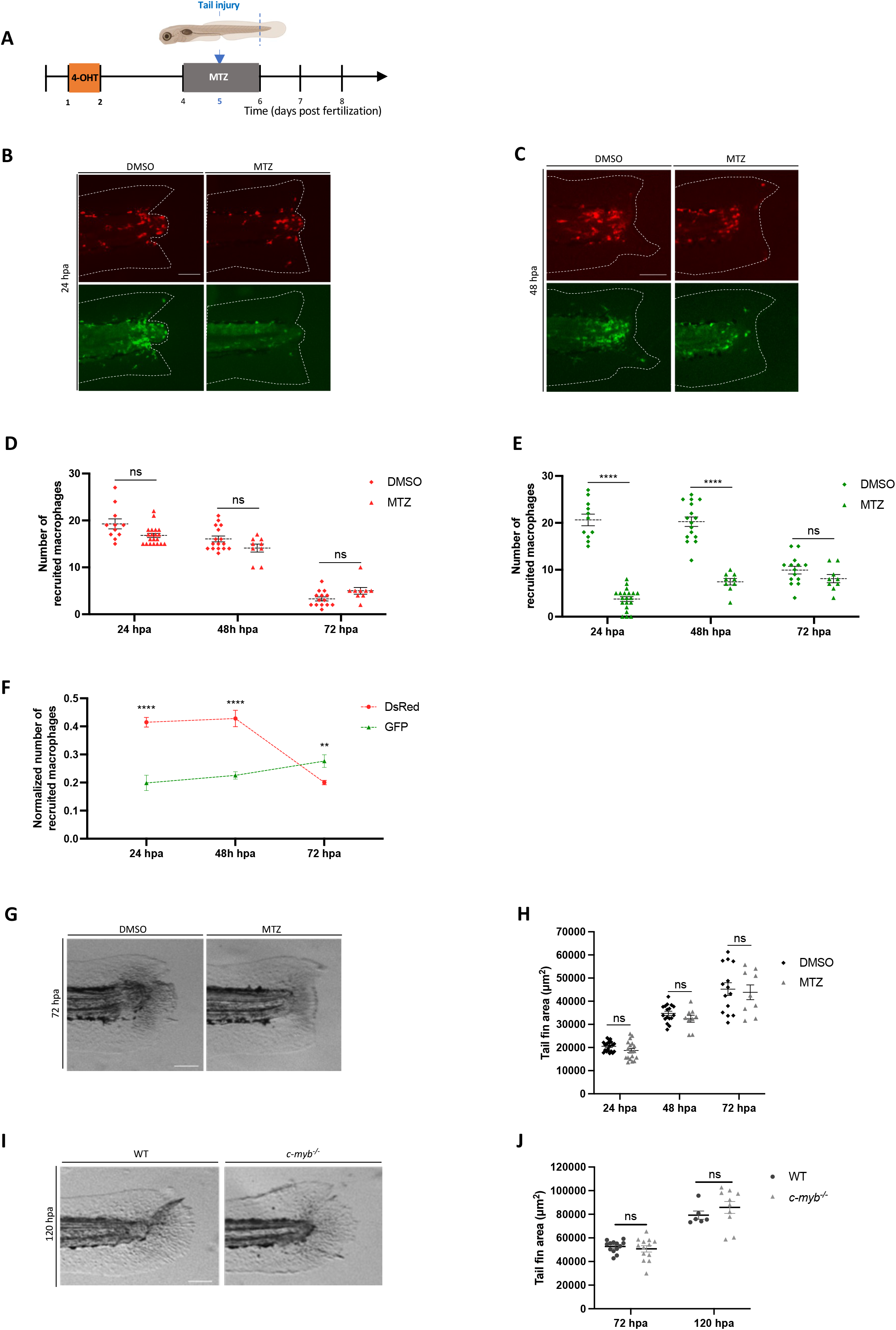
Selective ablation of definitive macrophages does not impair tail fin regeneration in early larvae. (A) Diagram showing the macrophage ablation and tail fin amputation strategy. (B-C) Switched *Tg(fli1a:creERT2;mpeg1:Switch)* larvae were treated with DMSO as a control, or metronidazole (MTZ) to ablate definitive macrophages. Treatments were performed from 4 to 6 dpf and tail fins were amputated at 5 dpf. Representative images are shown 24 hpa in (B) and 48 hpa in (C). Scale bars: 100 µm. (D) Quantification of DsRed+ macrophages in the tail region at 24, 48 and 72 hpa in larvae treated at 4 dpf with either DMSO or MTZ for 48 hours (24 hpa: DMSO=11, MTZ =20; 48 hpa: DMSO n=16, MTZ n=9; 72 hpa: DMSO n=14, MTZ n=9). Mean ± SEM ofthe DsRed^+^ macrophage number is shown. ns, p>0.05. (E) Quantification of GFP+ macrophages in the tail region at 24, 48 and 72 hpa in larvae treated at 4 dpf with either DMSO or MTZ for 48 hours (24 hpa: DMSO=11, MTZ =20; 48 hpa: DMSO n=16, MTZ n=9; 72 hpa: DMSO n=14, MTZ n=9). Mean ± SEM of the GFP^+^ macrophages is shown. ns, p>0.05; ^∗∗∗∗^p ≤ 0.0001. (F) Quantification of DsRed+ and GFP+ macrophages at the site of injury at 24, 48 and 72 hpa. Quantification was normalized by the number of total macrophages in the tail of the respective larvae (the sum of peripheral, CHT and recruited macrophages of distinct origins). ^∗∗^p ≤ 0.01; ^∗∗∗∗^p ≤ 0.0001. (G) Representative images of regenerating tail fins of larvae at 72 hpa. Larvae were treated with either DMSO or MTZ. Scale bars: 100 µm. (H) Tail fin area quantification of regenerating tail fins at 24, 48 and 72 hpa in larvae treated with eitherDMSO or MTZ (24 hpa: DMSO n=21, MTZ n=20; 48 hpa: DMSO n=17, MTZ n=9; 72 hpa: DMSO n=14, MTZ n=9). Mean ± SEM of the tail fin area is shown. ns, p>0.05. (I) Representative images of regenerating tail fins of *cmyb*^null^ and WT larvae at 120 hpa. Scale bars: 100 µm. (J) Tail fin area quantification of regenerating tail fins of *cmyb*^null^ and WT larvae at 72 hpa and 120 hpa (72 hpa: *cmyb*^null^ n=13, WT n=13; 120 hpa: *cmyb*^null^ n=10, WT n=6). Mean ± SEM of the tail fin area is shown. ns, p>0.05.

We next performed tail fin amputation in 5 dpf larvae that was treated with either DMSO or MTZ from 4-6 dpf to ensure ablation of definitive macrophages through the regeneration process. While primitive macrophage numbers and recruitment were not affected during the regeneration process, definitive macrophages were significantly reduced during the first 48 hpa but recovered by 72 hpa (Figure 4B-E and supplemental Figure 3E). Next, we analyzed macrophage recruitment to the site of injury in MTZ-treated larvae from 24 to 72 hpa. We normalized the number of recruited macrophages by the number of total macrophages in the tail at their respective timepoints and we observed a decrease in the number of recruited definitive macrophages during the first 48 hpa, indicating the efficiency of the MTZ-mediated selective ablation of definitive macrophages (Figure 4F). We then performed tail fin regeneration analysis and observed no differences in the regenerated tail fin area between larvae treated with either DMSO or MTZ (Figure 4G-H). Furthermore, cell proliferation was not altered in the regenerated tail of 72 hpa larvae with either DMSO or MTZ (Supplemental Figure 3F-G).

To determine if this observation could be replicated in the absence of definitive macrophages, we analyzed tail fin regeneration in *cmyb*-deficient zebrafish that lack HSCs, have impaired definitive hematopoiesis and have reduced numbers of definitive macrophages^7,33,34^. We found no differences in regeneration efficiency, measured by the tail fin area, between *cmyb*-deficient and wild type larvae (Figure 4I-J). Our results, thus, indicate that primitive macrophages recruitment to the damage site is not affected and tail fin regeneration is not impaired when definitive macrophages are depleted.

### Depletion of primitive macrophages impairs tail fin regeneration in early larvae

To characterize the role of primitive macrophages in tail fin regeneration during early larval developmental stages, we chemically depleted them at 48 hpf, before the maturation of their definitive counterparts as previously reported in zebrafish^35^ (Figure 5A). In line with previous studies^29,30,35^, intravenous injection of L-clodronate at 48 hpf efficiently depleted macrophages in 5 dpf zebrafish larvae (Figure 5B-C). Thus, we performed tail fin amputation and regeneration analysis in L-clodronate injected larvae and compared them with L-PBS injected controls (Figure 5D-E). We found fewer macrophages at the injury site in L-clodronate injected fish compared to L-PBS injected controls (Figure 5F), suggesting primitive macrophage ablation at 72 and 120 hpa. In contrast to the selective ablation of definitive macrophages (Figure 4), depletion of primitive macrophages impaired tail fin regeneration, as measured by the tail fin area, at 72 hpa and 120 hpa (Figure 5D-E). These observations suggest that primitive macrophage depletion leads to tail fin regeneration defects, highlighting the functional differences among macrophages based on their ontogeny. Overall, these results shed light on the critical role of early-recruited primitive macrophages during tail fin regeneration in early zebrafish larvae.

**Figure 5:**
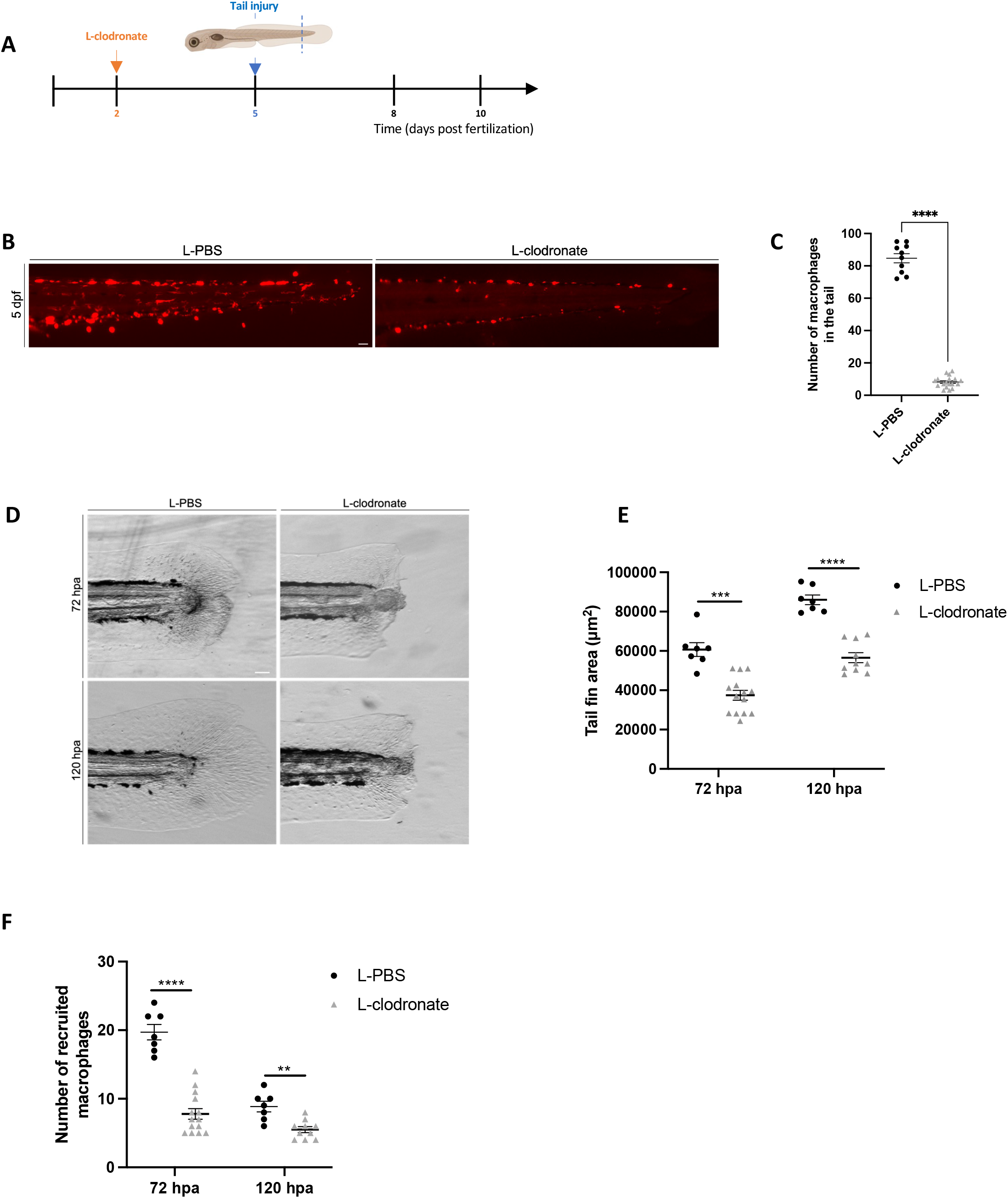
Depletion of primitive macrophages impairs tail fin regeneration in early larvae. (A) Diagram showing the macrophage depletion using L-clodronate injection and tail fin amputation plan. (B) *Tg(mpeg1:Switch)* larvae were injected at 48 hpf with L-PBS as a control, or L-clodronate (L-clo) to deplete primitive macrophages. Representative images are shown at 5 dpf. Scale bars: 100 µm. (C) Quantification of DsRed+ macrophages in the tail region at 5 dpf in larvae injected with either L-PBS (n=10) or L-clo (n=18). Mean ± SEM of the DsRed^+^ macrophages is shown. ^∗∗∗∗^p ≤ 0.0001. (D) Representative images of regenerating tail fins of larvae at 72 and 120 hpa. Larvae were injected with either L-PBS or L-clodronate. Scale bars: 100 µm. (E) Tail fin area quantification of regenerating tail fins at 72 and 120 hpa in larvae injected with either L-PBS or L-clodronate (72 hpa: L-PBS n=7, L-clo n=14; 72 hpa: L-PBS n=7, L-clo n=10). Mean ± SEM of the tail fin area is shown. ^∗∗∗^p ≤ 0.001; ^∗∗∗∗^p ≤ 0.0001. (F) Representative images of macrophages at the site of injury at 72 and 120 hpa in larvae injected with either L-PBS or L-clodronate. Scale bars: 100 µm. (G) Quantification of macrophages at the site of injury at 72 and 120 hpa (72 hpa: L-PBS=7, L-clo n=14; 120 hpa: L-PBS=7, L-clo n=10). Mean ± SEM of the macrophage number is shown. ^∗∗^p ≤ 0.01; ^∗∗∗∗^p ≤ 0.0001.

## Discussion

Two major waves of hematopoietic progenitors are generated during embryogenesis: the primitive and the definitive waves. Both waves generate erythromyeloid progenitors during embryogenesis, however, having a dual source of blood progenitors makes it difficult to assess accurately each wave contribution to blood lineages in homeostasis and upon tissue injury. Therefore, the lack of specific markers to identify different populations from the same blood lineage according to their origin and function has hindered the full understanding of immune cell ontogeny.

Here, we classified hematopoietic waves according to their site of origin and tracked them during homeostasis and upon recruitment to the site of injury. Our study shows that the hematopoietic system follows a layered strategy to provide a timely supply of innate immune cells and erythrocytes. Under steady-state conditions, we found that the contribution of definitive hematopoietic progenitors to embryonic erythromyeloid lineage is limited. This is of particular interest as none of the lineage tracing strategies performed in mice, despite their elegant designs, are specific to definitive hematopoietic waves. Our results also complement the recently reported observations that HSCs have a delayed contribution to the lymphomyeloid lineage during development ^10^.

In addition, as timely recruitment of macrophages to the damage site is important to ensure proper tail fin regeneration ^28–31^, we found that the primitive macrophages are the first responders after tail fin injury showed by their active phagocytic activity as early as 3 hpa and their earlier recruitment than their definitive counterparts. The difference in recruitment behaviour between primitive and definitive macrophages could be due to that the primitive macrophages are more sensitive to damage sensing and/or a consequence of the immaturity of definitive macrophages at early developmental stages. However, we delineated that definitive macrophages regenerate following their depletion but are not required for tail regeneration during early developmental stages. These intriguing results suggest that embryos and early larvae adopt a layered ontogeny of macrophages to ensure their abundance for a timely contribution to tissue regeneration. Further studies are needed to determine the molecular signatures of these ontogenetically distinct populations accounting for their different behaviors.

A significant discovery from our study highlights the key role of primitive macrophages in orchestrating tissue regeneration. Depletion of primitive macrophages during early developmental stages had a negative impact on the regeneration of tail fins. In contrast, the selective ablation of definitive macrophages did not hinder tail fin regeneration. This conclusion is supported by additional experiments, where we independently targeted either the definitive or primitive macrophage populations using two distinct methods and then analyzed the regrowth of tail fins in amputated larvae. These results underscore the remarkable capacity of primitive macrophages to facilitate timely tissue regeneration.

In conclusion, our study provides insights into the ontogeny of the erythromyeloid lineage during embryonic/early larval developmental stages. Our findings support the notion that embryos show a sequential emergence of hematopoietic waves to ensure the abundance of macrophages required for tissue homeostasis and regeneration during a crucial developmental time window. In line with our results, it has been recently reported in mice ^11,36–38^ and in zebrafish ^8,10^ that embryonic and adult HSCs do not give rise to several blood subtypes at steady state, upon ablation of mature blood cells, and in response to immune challenge. These observations indicate that ensuring a rapid response to stress to maintain tissue homeostasis in an HSC-independent manner might be a conserved mechanism between species throughout their life.

## Limitations of the study

*fli1a* is expressed in mature thrombocytes^39^ and is expressed also at 10 hpf in the lateral mesoderm^40^.Therefore, we used an inducible system to label all definitive hematopoietic progenitors as they emerge from hemogenic endothelium at 24 hpf. Using our labeling strategy, we could not distinguish distinct definitive hematopoietic progenitors as they all share the expression of *fli1a^+^* as they go through EHT^1,3,10^,therefore, better tools are required. Furthermore, in our study we were not able to selectively ablate the primitive macrophage population because of lacking a specific marker to the primitive hematopoietic progenitors and their progeny. In addition, we were unable to guarantee that all definitive hematopoietic progenitors were successfully labeled. According to the tracing result that ∼80% of thymocytes were labeled, there is still a remaining ∼20% of cells that likely derived from unlabeled definitive progenitors due to the incomplete *loxP* recombination.

## STAR★Methods

### Key resources table

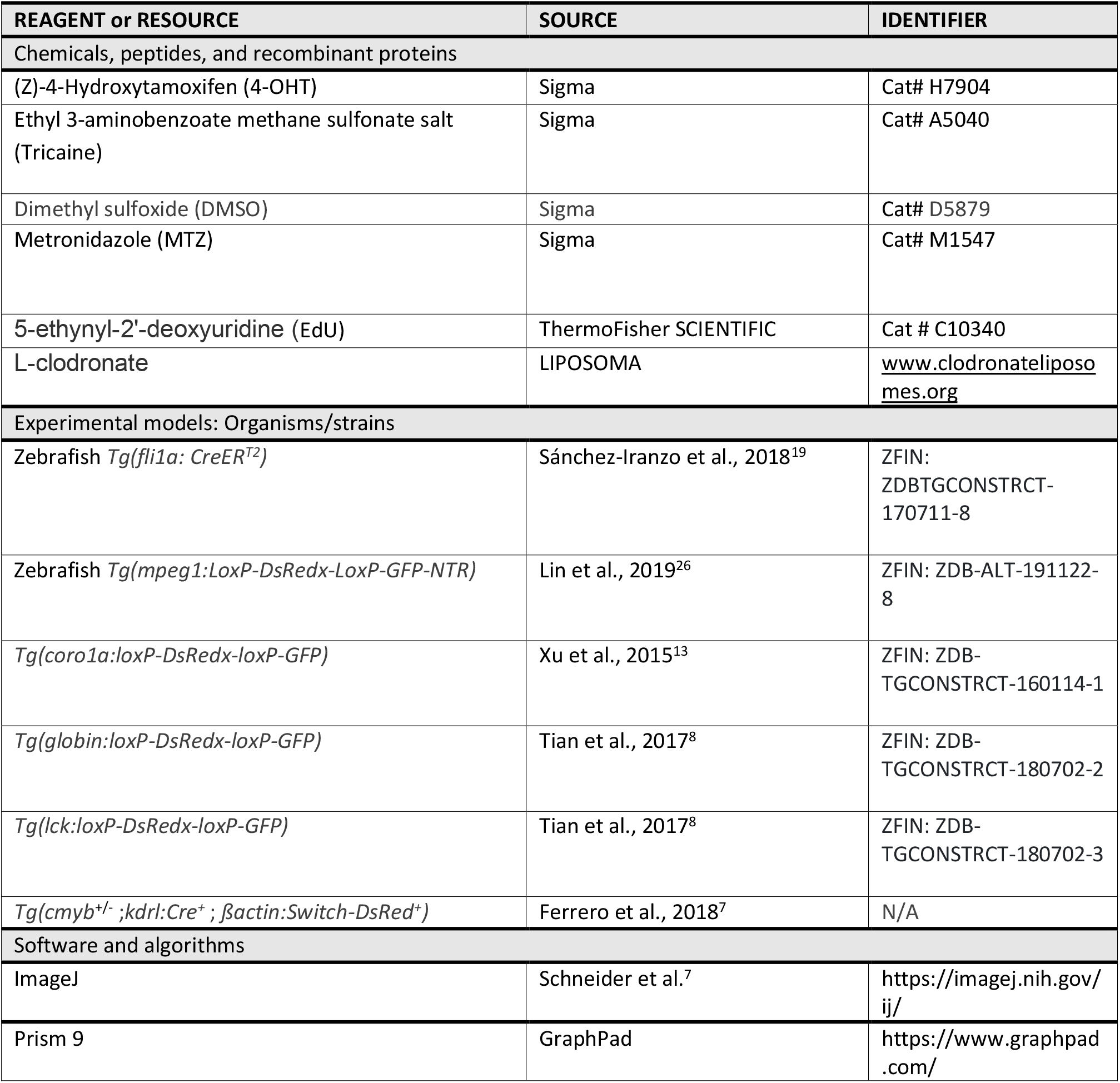

## Resource availability

### Lead contact

Further information and requests for scripts, resources, and reagents should be directed to and will be fulfilled by lead contact, Pedro P. Hernández.

## Materials and Data availability

This study did not generate new unique reagents. The data sets generated during the current study are available from the corresponding authors upon request.

## Experimental model and subject details

### Zebrafish

Embryonic and adult zebrafish (*Danio rerio*) were maintained at 28°C on a 14-hour light/10-hour dark cycle. The collected embryos were raised in fish water containing 0.01% <u>methylene blue</u> to prevent <u>fungal growth</u>. All fish are housed in the fish facility of our laboratory, which was built according to the local animal welfare standards. Animal care and use for this study were performed in accordance with the recommendations of the European Community (2010/63/UE) for the care and use of laboratory animals. Experimental procedures were specifically approved by the ethics committee of the Institut Curie CEEA-IC #118 (Authorization number - APAFiS#21197-2019062521156746-v2 - given by National Authority) in compliance with the international guidelines.”

### Transgenic lines

The following lines of the AB stain were used: *Tg(fli1a: CreER^T2^)*^19^*, Tg(mpeg1:LoxP-DsRedx-LoxP-GFP-NTR)*^26^,*Tg(coro1a:loxP-DsRedx-loxP-GFP)*^13^*, Tg(globin:loxP-DsRedx-loxP-GFP)* ^8^, *Tg(lck:loxP-DsRedx-loxP-GFP)* ^8^ *and Tg(cmyb*^+/-^; *kdrl:Cre^+^*; *ßactin:Switch-DsRed^+^)*^7^ *cmyb*^null^ embryos were identified at 5 dpf based on the absence of HSC-derived *DsRed^+^* thymocytes.

## Methods details

### Fluorescence Microscopy

Zebrafish embryos, larvae and adults were anesthetized with 0.01% tricaine (A5040, Sigma) and mounted in 2.5% 2.5% methylcellulose in 35-mm imaging dishes (MatTek) as described previously ^41^. Fluorescent imaging was performed with either Zeiss Axio Zoom V.16 upright microscope with an AxioCam HRm Zeiss camera and Zeiss Zen 3.3 software or with Leica thunder imaging system with Leica LAS-X software. Fluorescence was detected with dsRed and green fluorescent protein (GFP) filters.

### Image Analysis

All images were analyzed using FIJI software ^42^.

### CreER-loxP Cell Labelling (Lineage Tracing)

Zebrafish embryos at 24 hpf were treated with 5 μM 4-OHT (H7904, Sigma) for 24 hours. Controls were incubated in the equivalent amount of Ethanol solution during the same period. Light exposure was avoided by using foil to cover the plates as 4-OHT is light sensitive. After treatment, embryos were washed with fresh embryo medium and placed back in incubator. From 5 dpf, larvae were transferred to the fish facility nursery where they were kept and fed. Embryos were raised for further analysis at different developmental stages.

### Macrophage ablation and drug treatments

The *Tg(fli1CreER^T2^;mpeg1:loxP-DsRedx-loxP-GFP-NTR)* embryos were immersed in system water containing 10 mM metronidazole (M1547, Sigma) for 48 hours, which caused an acute depletion of GFP-NTR^+^ cells. The medium containing the MTZ was changed every 24 hours. Controls were incubated in the equivalent amount of DMSO solution during the same period. Light exposure was avoided by using foil to cover the plates as MTZ is light sensitive. Zebrafish were further analyzed by fluorescence imaging at different developmental time points. For primitive macrophage ablation using L-clodronate injection, the larvae were anesthetized in 0.016% Tricaine and microinjected with 5 nl of liposome encapsulated clodronate (www.clodronateliposomes.org) in the posterior caudal vein in the Urogenital Opening region at 48 hpf. Control embryos were similarly injected with liposome-PBS (L-PBS).

### Tail Fin Amputation

2 and 5 dpf larvae were anesthetized with 0.01% tricaine and amputated with a sterile scalpel. The amputation was performed by using the posterior section of the ventral pigmentation gap in the tail fin as a reference, and immediately after amputation larvae were incubated in either E3 medium, or in MTZ in case of macrophage ablation at 28°C.

### Quantification of Tail Fin Regeneration

At 24-, 48- and 72-hours post amputation or dpa (6,7- and 8-days post fertilization, respectively), larvae were mounted in 2.5% methylcellulose, and regenerating tail fins were imaged in bright field. Tail fin area was measured from the anterior section of the ventral tail fin gap to the end of the regenerating fin, as previously described^43^, using FiJi software. Tail fin areas were calculated and expressed in square micrometers (um^2^).

### Quantification of Tail Macrophages

Macrophages were quantified based on their location in the tail (periphery vs CHT) (Figure 2B) in non-amputated larvae. In amputated larvae, recruited macrophages to the site of injury were quantified. Normalization of recruited macrophage numbers was obtained by dividing the number of recruited macrophage subpopulation by their total number in the tail of the same larvae (periphery, CHT and recruited macrophages) ^29^. We have factored the labelling efficiency in our quantification data analysis in Figure 3 and Figure 4.

### Cell Proliferation analysis

For cell proliferation analysis, EdU analysis was performed in 72 hpa larvae previously fixed in 4% PFA and dehydrated with methanol, using Click-iT EdU cell proliferation kit for imaging, Alexa Fluor 647 dye. Stained larvae were imaged using fluorescence microscopy and analyzed using FIJI.

### Quantification and statistical analysis

Statistical analyses were performed by the GraphPad Prism software (Prism 9). All experiments with only two groups and one dependent variable were compared using an unpaired t-test with Welch’s correction. Two-way ANOVA with Sidak’s multiple comparison was used for this analysis. Statistical data show mean ± s.e.m. Each dot plot value represents an independent embryo, and every experiment was conducted three times independently. The corrected total cell fluorescence (CTCF) was calculated using the formula ‘Integrated density whole – (area whole embryo x mean fluorescence background)’. This formula is loosely based on a method described for calculating cell-fluorescence ^10^. To calculate the CTCF percentage for DsRed+ cells, the CTCF value of DsRed was divided by the total summation of CTCF values for both DsRed and GFP in each larvae. The same was done for calculating CTCF percentage for GFP+ cells.

## Acknowledgments

We thank the Developmental Biology Curie imaging facility (PICT-IBiSA@BDD, Paris, France, UMR3215/U934) members of the France-BioImaging national research infrastructure for their help and advice. We also thank the members of the animal facility of Institut Curie for zebrafish care. We also thank Yohanns Bellaiche and all the members of the Hernandez lab for thoughtful and valuable discussions. We are grateful to Nadia Mercader for providing the *Tg(fli1a:CreERT2)*, we also thank Zilong Wen for providing all the immune and blood lineage switch lines used in this study. This work was supported by the Institut Curie, INSERM, CNRS, the Ville de Paris emergence program (2020 DAE 78), FRM amorçage (AJE201905008718), ATIP-Avenir starting grant R21045DS and the Laboratoire d’Excellence (Labex) DEEP (ANR-11-LBX-0044, ANR-10-IDEX-0001-02 PSL), ECOS-ANID C22S01-220029. R.E. was supported by the Springboard postdoctoral fellowship from Institut Curie and Labex DEEP.

## Authors contribution

Conceptualization: R.E. and P.P.H.; Methodology and data collection: R.E., A.M., K.C., Y.S., P.D. and G.G.; Writing - Original Draft: R.E.; Writing - Review & Editing: R.E. and P.P.H.; Funding Acquisition: P.P.H.

## Conflict of Interest

The authors declare no competing interests.

---

**Sup. Fig. 1.**
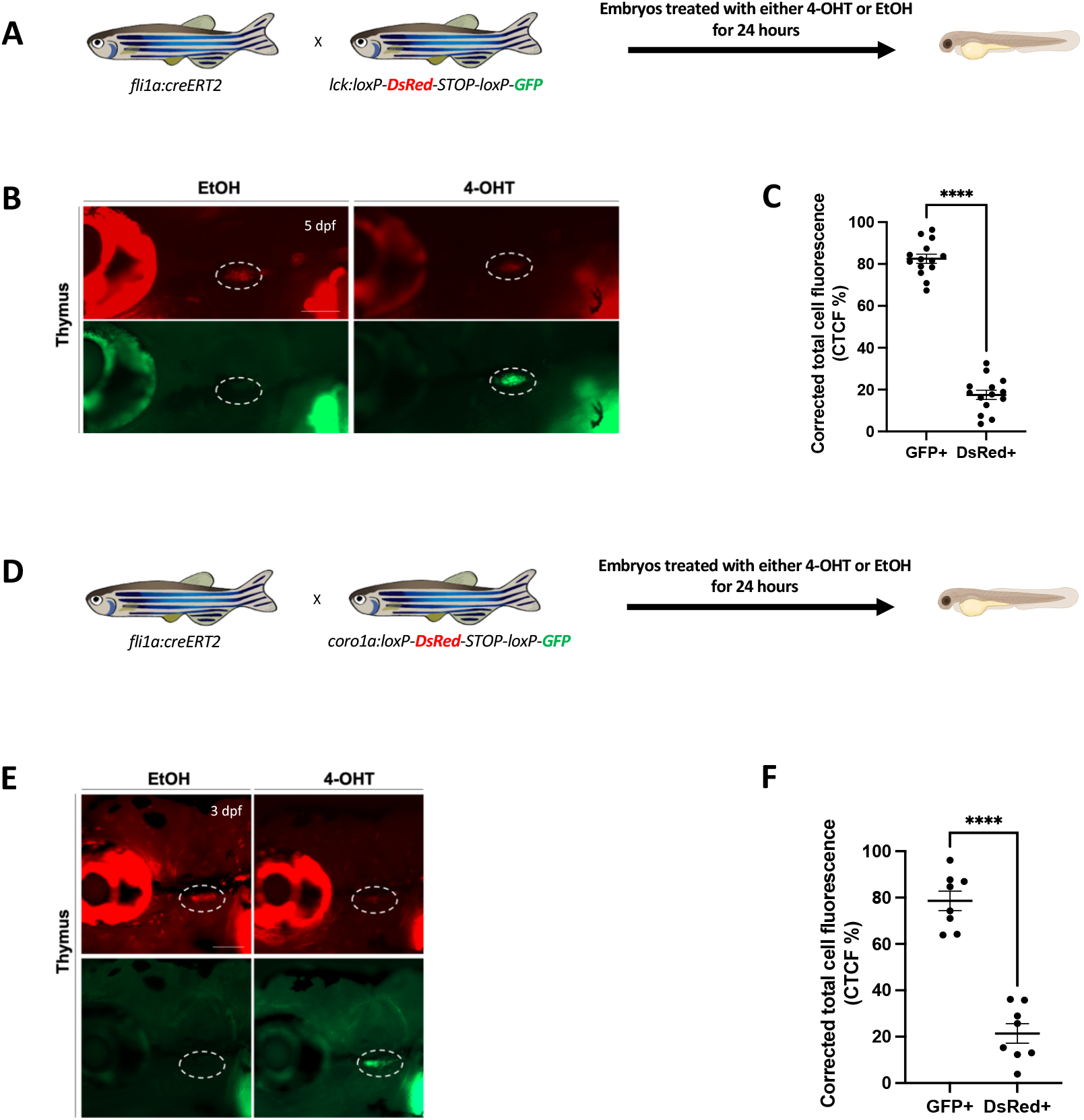
Labeling strategy and the labeling efficiency of using thymocytes as a positive read out with *lck*:switch and *coro1a*:switch transgenic lines. (A) Scheme of the 4-OHT-inducible *Tg(fli1a:creERT2;lck:Switch)* line used to assess the efficiency of the labelling strategy. (B) Fluorescent images of EtOH non-switched controls (left) and 4-OHT-induced (right) *Tg(fli1a:creERT2;lck:Switch)* thymocytes at 5 dpf larvae. Representative images are shown. Scale bars, 100 µm. (C) Quantification of DsRed and GFP fluorescence intensity percentage in the thymic region was measured at 5dpf (n=14). Mean ± SEM of the DsRed^+^ and GFP^+^ corrected total cell fluorescence (CTCF) percentage is shown. ****p≤ 0.0001 (D) Scheme of the 4-OHT-inducible *Tg(fli1a:creERT2; coro1a:Switch)* line used to assess the efficiency of the labelling strategy. (E) Fluorescent images of EtOH non-switched controls (left) and 4-OHT-induced (right) *Tg(fli1a:creERT2;coro1a:Switch)* thymocytes at 3 dpf embryos. Representative images are shown. Scale bars, 100 µm. (F) Quantification of DsRed and GFP fluorescence intensity percentage in the thymic region was measured at 3dpf (n=8). Mean ± SEM of the DsRed^+^ and GFP^+^ corrected total cell fluorescence (CTCF) percentage is shown. ****p≤ 0.0001

**Supp videos related to Fig. 1.**
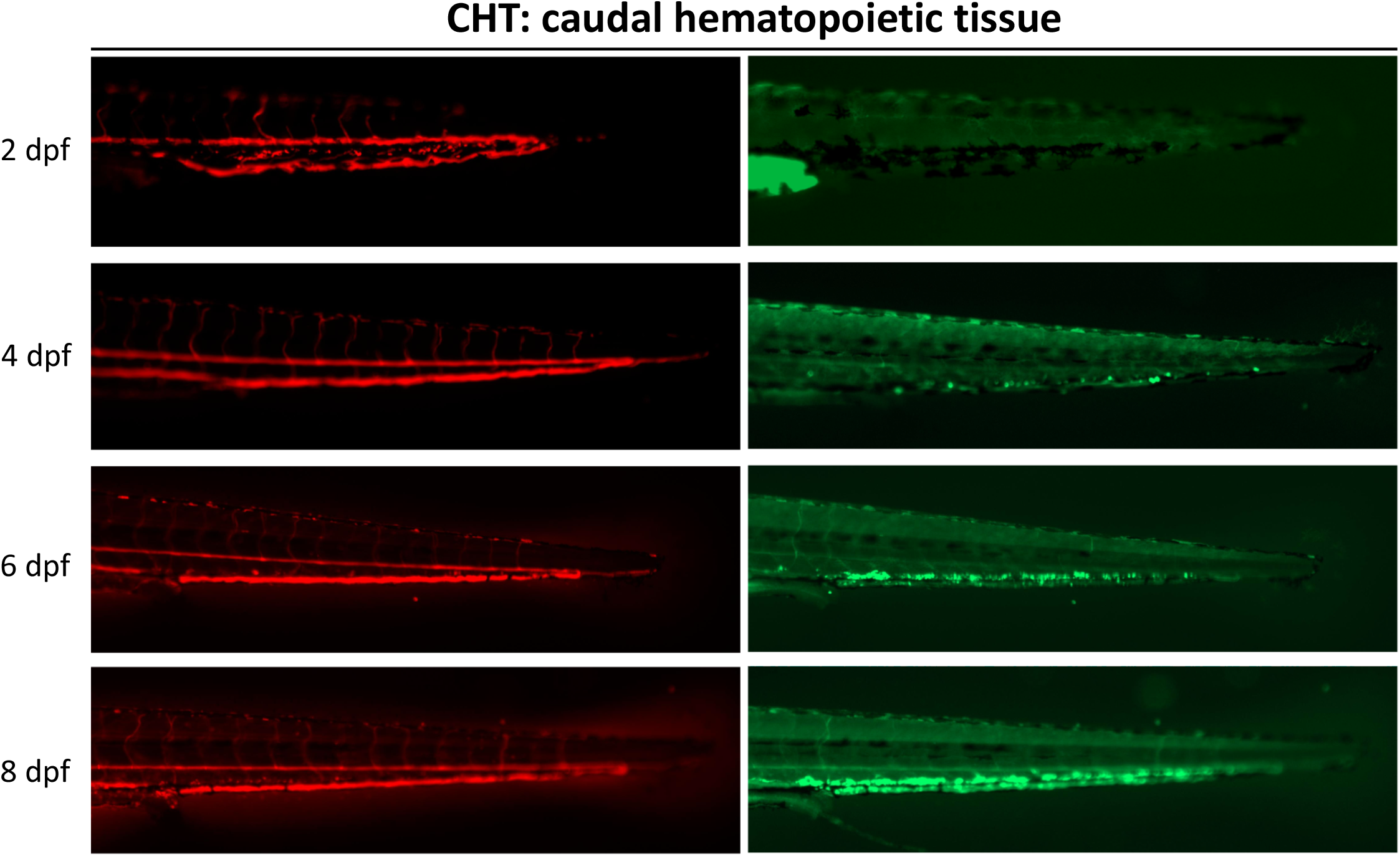
Fluorescence imaging of 4-OHT-induced *Tg(fli1a:creERT2;globin:Switch)* embryos and larvae over a time course of 2-8 dpf. Videos are assembled from different Z-stacks projections at each defined developmental stage.

**Supp Fig. 2.**
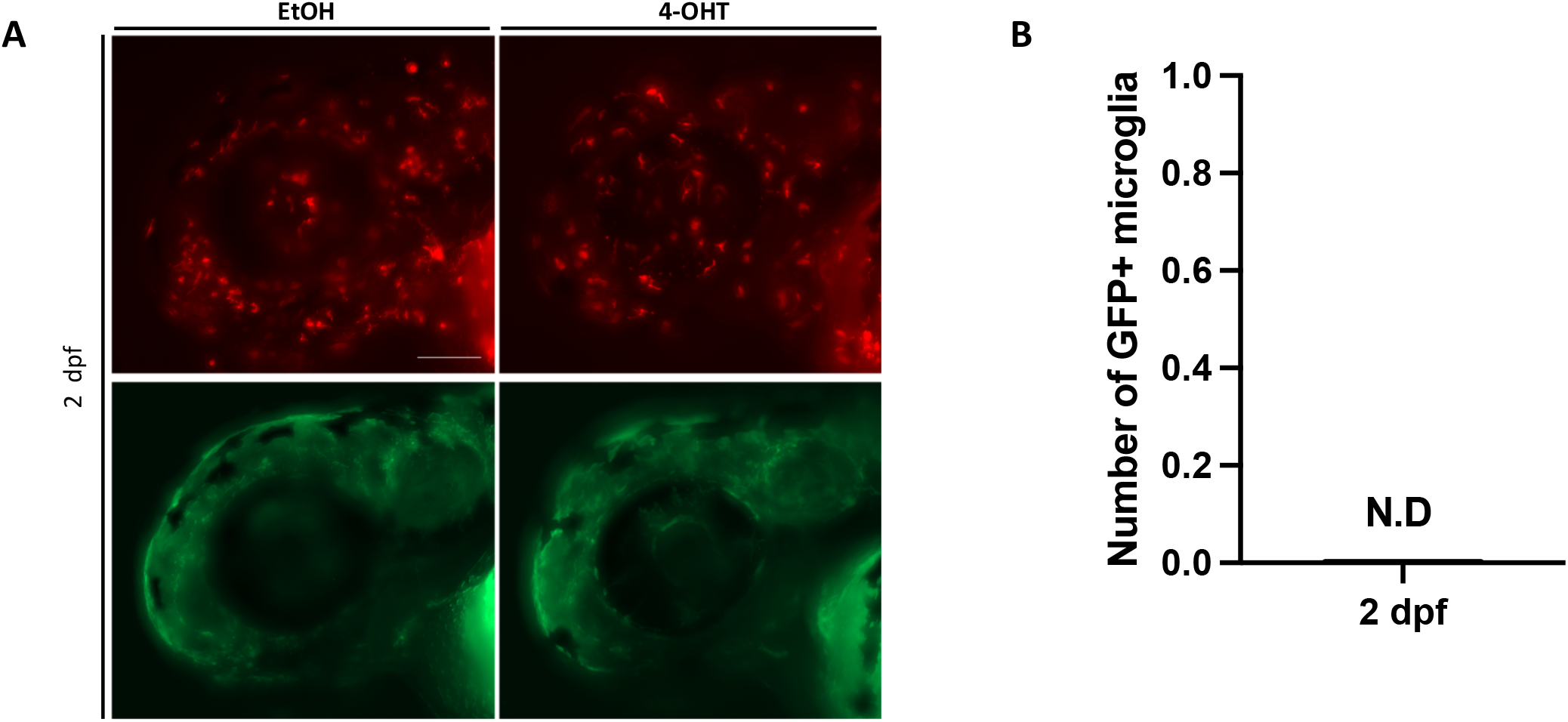
Quantification of GFP+ microglia in the head region at 2 dpf embryos. related to Fig.2. (A) Fluorescent images of EtOH non-switched controls (left) and 4-OHT-induced (right) *Tg(fli1a:creERT2;mpeg1:Switch)* head regions in the developing embryos at 2 dpf. Representative images are shown. Scale bar: 100µm. (B) Quantification of GFP+ macrophage number in the head at 2 dpf (n=6). Mean ± SEM of the GFP^+^ macrophage numbers are shown.

**Supp Fig 3. related to Fig. 3 and 4.**
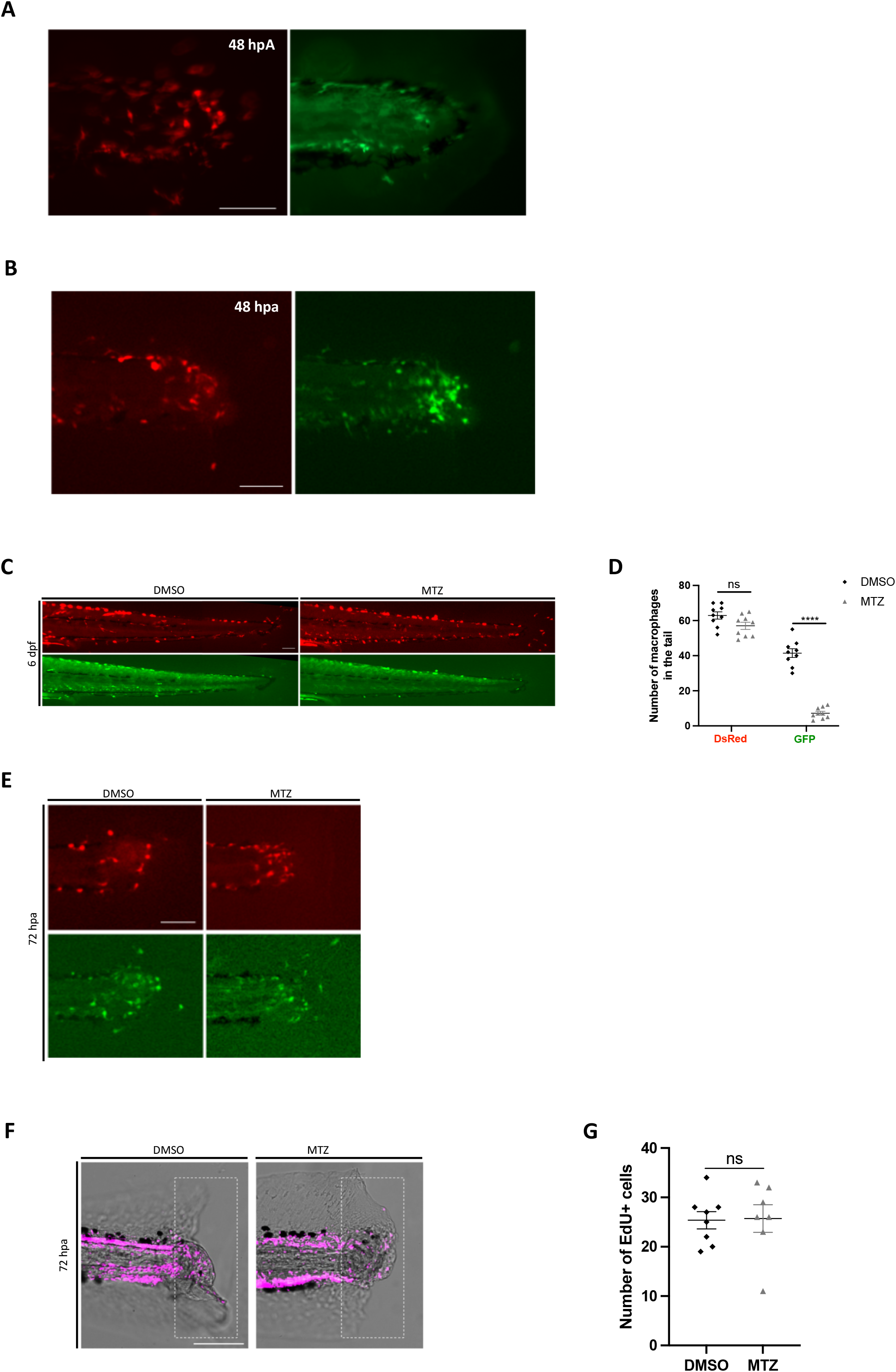
Ontogenetically distinct macrophages recruitment after tail injury at different developmental stages. (A) Tail fin of *Tg(fli1a:creERT2;mpeg1:Switch)* were amputated at 2 dpf and macrophage recruitment to the site of injury was analyzed. Representative images are shown at 48 hpa. Scale bar: 100µm. (B) Tail fin of *Tg(fli1a:creERT2;mpeg1:Switch)* were amputated at 5 dpf and macrophage recruitment to the site of injury was analyzed. Representative images are shown at 48 hpa. Scale bar: 100µm. (C) Switched *Tg(fli1a:creERT2;mpeg1:Switch)* larvae were treated with DMSO as a control, or metronidazole (MTZ) to ablate definitive macrophages. Treatments were performed from 4 to 6 dpf. Representative images are shown at 6 dpf. Scale bars: 100 µm. (D) Quantification of DsRed and GFP macrophage number in the tail region 48 hours after DMSO (n=9) or MTZ treatments (n=9). Mean ± SEM of the DsRed^+^ and GFP^+^ macrophage numbers are shown. ^∗∗∗∗^p ≤ 0.0001. (E) Switched *Tg(fli1a:creERT2;mpeg1:Switch)* larvae were treated with DMSO or MTZ. Treatments were performed from 4 to 6 dpf and tail fins were amputated at 5 dpf. Representative images are shown 72 hpa. Scale bars: 100 µm. (F) Cell proliferation was measured by EdU assay at 72 hpa. Red dots represent EdU^+^ cells. Representative images are shown at 72 hpa of larvae treated with DMSO as a control, or metronidazole (MTZ) to ablate definitive macrophages. Scale bar: 100µm. (G) Numbers of EdU^+^ cells in switched *Tg(fli1a:creERT2;mpeg1:Switch)* larvae that were treated with DMSO (n=8) or MTZ (n=7). Unpaired t-test with Welch’s correction was used for the analysis.

## Notes

### Competing Interest Statement

The authors have declared no competing interest.

### Summary of Updates

To ablate primitive macrophages to assess their contribution to tail regeneration in zebrafish larvae.

